# Anaerobic gut fungi *Caecomyces communis, Neocallimastix frontalis* and *Piromyces spp. nov.*, have distinct effects on plant fibres during digestion

**DOI:** 10.1101/2025.09.21.677623

**Authors:** Shuqi Shen, Jessica L. Matthews, Siwei Li, Jolanda M van Munster

**Author notes:** For correspondence: Jolanda van Munster.

## Abstract

Anaerobic gut fungi are the first colonizers of plant material that enters the digestive system of ruminants. However, it is unclear how different fungal species contribute to the ability of the rumen microbiome to convert feed to nutrients. Here we isolated three species of anaerobic fungi, including a novel *Piromyces* species. We investigated if these species have distinct roles in the digestion of fibrous feed components, via assessment of changes caused to plant material itself. We found *Neocallimastix frontalis* isolate CoB3 and *Piromyces* isolate SHC digested plant materials more effectively than *Caecomyces communis* isolate SHB. The three fungi had distinct effects on feed composition. *N. frontalis* CoB3 degraded hemicelluloses and cellulose to a similar extent, *Piromyces spp. SHC* preferentially degraded hemicellulose, while *C. communis SHB* preference depended on the substrate. From the panel of monosaccharides that may result from such degradative activity, all fungi consumed only glucose, suggesting involvement of mechanisms more complex than only fungal carbon source usage. Overall, this indicates that each of these fungal species have distinct roles in the degradation of plant material, and different niches in the rumen. Exploring these roles creates functional understanding of the rumen microbiome, critical for developing more sustainable agriculture.

## Introduction

Feed digestion is critical to ruminant performance in farming [1]. Activity of the microbial community in the digestive tract of ruminants, such as cows and sheep, enables conversion of the plant cell walls of the feed to nutrients such as short chain fatty acids. This enables ruminants to convert feed and forages that are inedible by humans, to high quality protein milk and meat. Ruminant farming thereby contributes to meeting the dietary requirements of the growing world population, especially in low- and middle-income countries [2], [3]. However, sustainable farming methods are required to ensure animal performance and mitigate environmental effects [4], [5]. Metabolic activity of the rumen microbiome produces the greenhouse gas methane, whereby ruminant farming results in 14% of anthropogenic methane emissions [6] or ~ 90% of livestock-associated methane emissions, which primarily (56 - 75 %) is caused by enteric fermentation [7]. Methane emission also results in 3 - 10% of feed energy loss [8], [9], exemplifying that the activity of the rumen microbiome affects feed efficiency. Microbiome manipulation is therefore a promising avenue to address sustainability challenges in ruminant farming, but this requires more detailed understanding of microbiome function [10], in particular of that of eukaryote members.

Anaerobic gut fungi (AGF), phylum *Neocallimastigomycota*, are the fungi central to fibre degradation in the herbivore gut microbiome. They are found in ruminants as part of the rumen microbiome, but also in the digestive system of hindgut fermenters such as horses and in other herbivores such as tortoises [11], [12], [13]. AGF are considered to be the first colonisers of plant material that is ingested by their host [14]. The fungi break down this plant material using the highest diversity of degradative carbohydrate active enzymes identified in any fungal genome sequenced to date [15]. These enzymes can be arranged in multi-enzyme complexes similar to bacterial cellulosomes [16]. The fungal cellular structures penetrate the plant material [17], [18], [19] and are thought to thereby further aid degradation, opening up the structure to increase access for enzymes and for degradative bacteria (Gruninger et al., 2014). AGF are also able to digest various lignin-containing plant tissues, releasing lignin-derived compounds in the process [21], [22]. This is of interest in agriculture as well as biotechnology, as lignin hinders effective plant cell wall degradation by most other microbes and their enzymes.

The activities of AGF contribute considerably to the plant-degradative activity of the rumen microbiome. In *in vitro* cultures, activity of AGF in enrichment cultures derived from rumen fluid increased digestibility of the fibrous plant materials by 26-53% [23] and of legume and grasses by 13-16% [24], [25], when compared to bacterial enrichment cultures. Their strong fibre degradative activity translates to activity *in vivo*, where AGF, either native or used as direct-fed microbial supplement, increase feed digestion, weight gain and milk production *in vivo* in ruminants, as reviewed by [20]. For a step change towards fully understanding activity of the ruminant microbiome, it is thus essential to improve understanding of the roles of fibre-degrading AGF.

Herbivores typically harbour a large diversity of AGF in their digestive system [11]. This diversity suggests that these fungi have distinct roles and niches in the environment of the digestive tract. Indeed, the composition of the AGF community in the gut system varies according to conditions and specific properties of their host animals, such as the animal physiology, including digestive tract properties [11], [12], [13], life stage [26], [27] and the host’s diet [13], [28], [29], [30]. While all AGF fungi are recognised as being involved in fibre degradation, it is not well known what the exact roles of this broad array of AGF species are. We consider that one of the factors involved in the evolution of these different AGF genera or species is the ability to effectively degrade different plant cell wall components or structures in animal diets, providing them with different niches in plant fibre digestion. We thus hypothesize AGF species may have distinct abilities to degrade plant biomass, and have different effects on its structure and composition during its degradation.

Here we investigate the degradative effects of three genera of anaerobic gut fungi that are common in agriculturally important ruminants such as cattle and sheep. We assessed the biochemical effects of their degradative activity on the composition of distinct fibrous plant materials that are commonly part of ruminant diets, more specifically wheat straw and timothy grass hay. In doing so, we provide new insight in the roles of these fungi in plant biomass degradation, demonstrating that the investigated fungi have distinct effect on the plant material that they degrade, which is suggesting that the AGF may indeed have different niches in the rumen. Via this work we also provide a basis for further exploration of the underpinning mechanisms of these AGF for lignocellulose degradation and a more in-depth elucidation of their functional role in the rumen microbiome.

## 2. Material and methods

### 2.1 Fungal cultivation conditions

Fungal isolates were routinely maintained by twice-weekly passaging of 5% (v/v) of the cultures into fresh medium C, modified from [31] to replace bactocasitone with tryptone. As carbon source, 1% w/v wheat straw milled to 0.5 mm was added, and cultures were performed under CO_2_ atmosphere in Hungate tubes via stationary incubation at 39°C. To prevent bacterial contamination, the medium was routinely supplemented with 50 µg ml^−1^ chloramphenicol. For fungal isolations, roll tubes contained medium C with 18 gr l^−1^ agar, and 5 gr l^−1^ glucose as well as 5 gr l^−1^ cellobiose as carbon sources. Where required, fungal growth was monitored via fermentation gas production, a proxy for fungal growth recognised as accurate in the field [32], [33]. Fermentation gas pressure was measured via a Vernier Labquest 2 equipped with pressure sensor. Culture pH was measured directly after opening of the culture vessel, using an Orion Star A111 benchtop pH meter with a Fisherbrand FB68801 semi-micro pH probe.

### 2.2 Isolation of anaerobic gut fungi

Fresh faecal samples were collected from a pasture with grazing black-face sheep, and transported to the laboratory for processing within an hour of collection. Samples were dispersed in anaerobic water, and 100-fold dilutions were generated that were immediately used to inoculate Hungate tubes with medium C containing 1% w/v of lignocellulose substrate (40% hay, 60% straw) milled to 0.5 mm. Separately, bovine rumen fluid, obtained from a rumen-fistulated cow, was used to inoculate medium C, which was used to generate a 20-fold serial dilution series in Hungate tubes with medium C containing 1% lignocellulose substrate (hay (20%), barley (30%), and wheat straw (50%). To inhibit growth of bacteria and archaea, antibiotics were added (100 µg ml^−1^ penicillin, 50 µg ml^−1^ kanamycin, 50 µg ml^−1^ chloramphenicol and 100 µg ml^−1^ streptomycin) to the enrichment cultures. After incubation for up to two weeks or until fungal growth was observed via floating biomass and/or gas production, cultures were propagated every 4 days. Axenic cultures were subsequently obtained by at least three rounds of purification on solid media in roll tubes alternated with propagation in liquid medium. Purification was continued until axenic cultures were established.

### 2.3 Molecular characterisation and phylogenetic analysis

Fungal biomass, derived from a 40 ml culture grown for 5 days with glucose as carbon source, was dried by lyophilisation and disrupted using a pestle and mortar. Genomic DNA was extracted using QIAamp Fast DNA Stool mini kit (Qiagen) according to the manufacturer’s instructions. Obtained DNA was used as template for amplification of the internal transcribed spacer (ITS region) of the rDNA by PCR using the primers ITS1 (5'-TCCGTGGTGAACCTGCGG–3') and ITS4 (5'-TCCTCCGCTTATTGATATGC–3') [34]. The D1-D2 regions of 28S rDNA was amplified using primer pair NL1 (5′-GCATATCAATAAGCGGAGGAAAAG-3′) and NL4 (5′-GGTCCGTGTTTCAAGACGG-3′) [35]. PCR conditions were as described by [36] using GoTaq Green Master Mix (Promega). Amplified products were purified using Wizard SV Gel and PCR Clean-up System (Promega) following the manufacturer’s directions, and cloned into the pGem-T Easy vector (Promega). The ITS or rRNA gene fragment sequences were subsequently obtained via Sanger sequencing of clones. Obtained sequences were trimmed to remove the backbone vector sequence. Sequences representing different AGF genera and species, provided in supplementary file 1 (LSU) and supplementary file 2 (ITS), were derived from [37] and the AF_Full_Region database version 2.0 as obtained from the Anaerobic Fungi Network (www.anaerobicfungi.org). Sequence alignments were performed in Clustal Omega [38], using *Gonopodya prolifera* JN874506 as outgroup for LSU sequences [39]. Alignments were trimmed in BioEdit 7.7.1 and phylogenetic analysis was performed via maximum likelihood test with 1000 bootstraps in IQ-TREE [40]. Tree files were visualised in MEGA11 [41]. Sequence identities were calculated as recommended [42], using relative blastn with the following parameters: match score 2; mismatch score −3; gap existence cost 5; gap extension cost 2.

### 2.4 Effects of fungal activity on plant material

For fibre analysis experiments, timothy grass hay was obtained commercially, while straw from wheat variety Barrel was obtained from Drumalbin farm, Lanark, Scotland. Both straw and hay were milled using a Retsch - SM100 cutting mill with a 2 mm screen mesh. Fungal cultures were performed in either 200 ml medium C in 250 ml Duran GL45 pressure plus bottles or 100 ml medium in 125 ml bottles, sealed with bromobutyl rubber septa. Cultures contained 50 µg ml^−1^ chloramphenicol and 1% w/v plant material, either wheat straw or timothy hay milled to <2 mm. Inoculations were performed via transfer of 5% v/v (10 ml) of 3 d old 6 ml starter cultures grown on wheat straw, into the experimental culture.

Cultures were sacrificed at set time points and solids were collected by centrifugation at 4500 rpm for 12 minutes, heat inactivated at 95 °C for 15 minutes, cooled and stored at −20 °C until analysis. Samples were lyophilised for 48h and weighed immediately after. For fibre analysis, 0.5 gr of culture solids was sealed in a F57 filter bag (ANKOM technology), and Neutral detergent fibre (NDF) and acid detergent fibre (ADF) were determined according to [43] employing the automated ANKOM 200 Fibre Analyser. Heat-stable bacterial alpha-amylase enzyme (FAA, ANKOM technology) was used for the determination of NDF following the manufacturer’s instructions. Hemicellulose content was calculated as NDF – ADF. Results were analysed via either ANOVA followed by post-hoc Tukey’s HSD test or a Kruskal Wallis test.

### 2.6 Growth on soluble sugars

For testing of fungal growth on monosaccharides, cultures were grown in Hungate tubes containing 6 ml Medium C as described above, whereby different sugars were added as carbon source. Sugars were added as filter-sterilised solution after autoclaving, to reach a 5 gr l^−1^ concentration.

## 3. Results

### 3.1 Isolation and identification of three species of anaerobic fungi, including a so far undescribed *Piromyces species*

With the aim of obtaining distinct species of anaerobic fungi that are common in agriculturally important ruminants, we performed fungal enrichment cultures via inoculation of either bovine rumen fluid or sheep stool samples in anaerobic media containing milled plant material as carbon source. From these cultures, a panel of fungal strains was isolated via sequential colony picking in roll tubes and further cultivation in liquid media. Assessment of fungal morphology by microscopy indicated that isolate SHB (Fig. 1A) had bulbous cells, while isolates SHC (Fig. 1B, Fig. S2) and CoB3 (Fig. 1C) had a filamentous rhizoid system, with either short and heavily branched rhizoids (Fig. 1B) or long rhizoids (Fig. 1C). Phylogenetic analysis of the ribosomal ITS (Fig. S1) and 28S rRNA gene (LSU) sequences (Fig. 1D) of these isolates indicated that species from three genera were obtained. Isolate CoB3 represents a *Neocallimastix frontalis* isolate, as indicated by strong clustering of ITS and LSU sequences with other isolates of species *Neocallimastix frontalis*, and, more remotely with other species in the *Neocallimastix* genus. The LSU sequence of isolate SHB has 98.0-100% identity to *C. communis* and *C. communis var. churrovis* respectively, and clustered with *C. communis* sequences, confirming that SHB is an isolate of species *C. communis*. The ITS sequences of *C. communis* isolate SHB and *C. communis var. churrovis* varied by only 1.3 - 2.8 %, confirming close similarity of these species.

**Figure 1.**
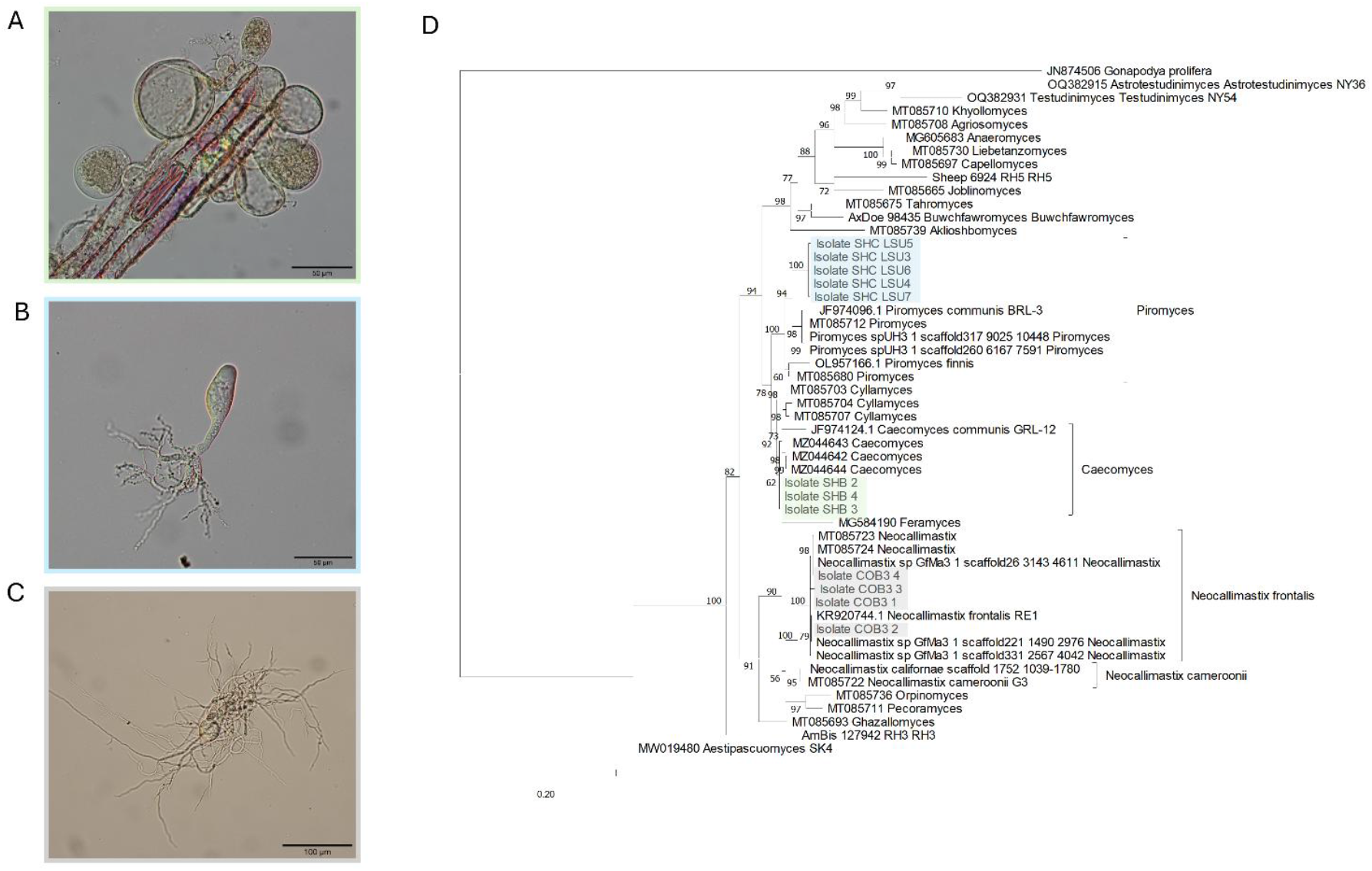
Isolation and identification of anaerobic gut fungi. a) *Caecomyces communis* isolate SHB grown for 10 days on wheat straw as carbon source, and b) *Piromyces spp* isolate SHC, c) *Neocallimastix frontalis* isolate CoB3, both grown 2 days on glucose. d) Maximum likelihood phylogenetic tree based on D1-D2 region of the LSU rDNA gene sequences of isolates described in this study. Isolates sequences are highlighted in green, blue, and grey, corresponding to the coloured frames around images in A-C.

The LSU sequences from isolate SHC clustered with high (100%) bootstrap support, with sequences of the *Piromyces* genus, closest to the proposed *P. indianae* spUH3 (96.8-97.0% sequence identity) and *P. finnis* (96 – 96.3% identity). Within isolate LSU variability was 0 - 0.26% over 5 clones. ITS1 sequence variation between the 4 obtained clones of isolate SHC ranged from 0 - 0.75%. The SHC ITS1 sequence identity to that of other *Piromyces* species was low, with 81.4 - 82.1% identity to *P. finnis*, 88.5 - 88.8 % to *P. indianae* SpUH3, 86.4 - 87.2 % to *P. communis* type strain P and 91.4 - 92.3% to *Piromyces communis* Jen1. This indicates that based on sequence data, isolate SHC represents a so far unidentified *Piromyces* species.

#### *Description of Piromyces spp. SHC* sp. nov

*Typification:* The holotype shown in Fig. 1C, is derived from the UK, Scotland, Edinburgh (55.95 N, −3.18 W), a 2-day old culture of strain SHC grown on media C containing glucose, originally isolated from fresh faeces from a sheep, January 2021. Ex-type strain: SHC. GenBank PX270344-PX270347 (ITS1-18S-ITS2) and PX270330-PX270334 (D1/D2 LSU).

An anaerobic fungus, forming small circular colonies on solid media in roll tubes, that are light brown with a dense, darker centre. Liquid growth is as a thin biofilm, with heavy attachment to container glass surfaces. Zoospores are spherical, 5 - 7 µm in diameter, monoflagellated with flagella 20 - 25 µm in length. Thalli are monocentric, with sporangia shaped irregularly with lateral bulges, triangular, heart-shaped or fusiform, 35 - 90 µm in diameter. Rhizoidal growth pattern is filamentous, with short, narrow and heavily branched hyphae.

### 3.2 Fungal growth on lignocellulose

In their natural environment of the rumen, AGF contribute to digestion of complex plant cell wall structures. We aimed to assess the ability and efficiency of the three obtained AGF isolates in fermenting wheat straw or timothy grass hay, which are commonly used as forage and roughage in ruminant diets. Wheat straw is also an abundant agricultural waste used as feedstock for renewables-based biotechnology.

A pilot experiment was performed, cultivating *C. communis* SHB and *P. spp* SHC on timothy grass hay (Fig. 2A), to determine time points along the growth curve for analysis. Accumulation of fermentation gas was used to quantify fungal growth, a method routinely used in the field as a direct relation between gas production and fungal growth has previously been demonstrated [32], [33]. From these growth curves, timepoints at day 2, day 3 and day 5 were selected to correspond with cultures in mid exponential, late exponential and start of the stationary growth phase respectively.

**Figure 2.**
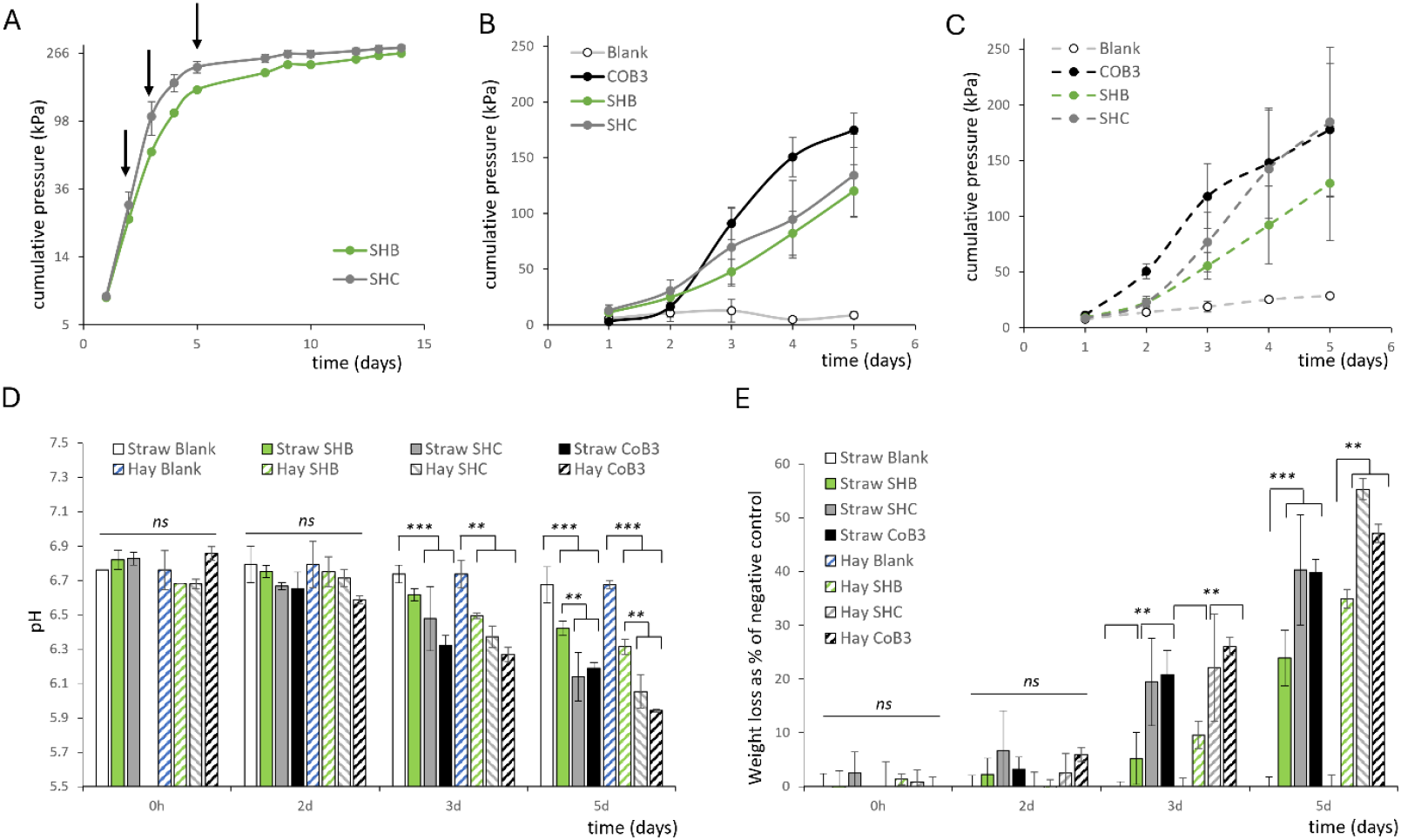
Growth of AGF on lignocellulose. a) Pilot growth curve of *P. spp* SHC (n=2) and *C. communis* SHB (n=1) on timothy hay, used to determine time points (arrows) in early, mid, and late exponential growth. Growth of 3 AGF species on wheat straw (b) and timothy hay (c) as detected by fermentation gas accumulation. Data shown is the mean ± stdev (n≥2), the blank did not contain fungal inoculant. d) pH of culture supernatant at day 5. e) Lignocellulose fermentation of wheat straw and timothy hay, expressed as consumption of culture solids compared to that in the negative control.

All three fungal isolates were subsequently grown on timothy grass hay or wheat straw as carbon source, with cultures sacrificed at day 2, 3 and 5 for collection of residual solids and culture filtrates. Growth, based on accumulation of fermentation gas, was comparable on both wheat straw (Fig. 2B) and timothy hay (Fig. 2C), and in line with results from the pilot experiment, growth curves displayed exponential growth phases around day 2. Overall, *N. frontalis* CoB3 accumulated ~ 1.4-fold more fermentation gas compared to *C. communis* SHB (Fig. 2B,C), and CoB3 had a higher specific growth rate on wheat straw (1.28 ± 0.07 µ day^−1^ for CoB3 compared to 0.75 ± 0.17 µ day^−1^ for SHB), as well as on timothy hay (1.49 ± 0.13 µ day^−1^ for CoB3, compared to 0.90 ± 0.10 on hay µ day^−1^ for SHB). *P. spp* SHC had both an intermediate gas accumulation and growth rates of 0.86 ± 0.25 µ day^−1^ on straw and 1.11 ± 0.19 µ day^−1^ on hay.

Analysed across both substrates, the pH in cultures of all fungi decreases significantly (*p*<0.003) in the day 3 and day 5 timepoints, compared to the corresponding negative controls (Fig. 2D). This is consistent with the ability of anaerobic fungi to produce organic acids such as acetate, formate and lactic acid during fermentation, accumulation of which would reduce culture pH. Overall, the final reduction in pH was significantly larger (*p*<0.003) for *N. frontalis* CoB3 and *P. spp* SHC than for *C. communis* SHB, further confirming differences in fermentation between *C. communis* SHB and both other fungal species.

To assess the substrate degradation efficiency of the three AGF, the dry weight of residual solids in the culture was measured and percentage of lost solids calculated. A reduction in dry weight of solids recovered from the cultures was only detected on day 5 of cultivation in cultures of SHB, while for SHC and CoB3 reductions in the obtained solids was observed from day 3 onwards (Fig. 2E). After 5 days of fermentation, *P. spp* SHC and *N. frontalis* CoB3 had solubilised more material than *C. communis* SHB, reducing the amount of wheat straw with 40 ± 10 % and 40 ± 2.5 % vs 24 ± 5.2 % respectively, and timothy hay with 55 ± 2.0 % and 47 ± 1.7 % vs 35 ± 1.8 % respectively. In the not-inoculated controls there was no difference in the weight of culture solids over the incubation time.

These results indicate that *P. spp* SHC and *N. frontalis* CoB3 degrade the tested plant materials both quicker and to a larger degree, and therefore overall more effectively, than *C. communis* SHB. Both pH and fermentation efficiency measures suggest that the timothy hay is more effectively fermented than wheat straw by all three fungi.

### 3.3 Distinct effects of fungal digestive activity on lignocellulose composition

To assess the effects of activity of the three fungal species on the composition of the plant material, fibre analysis was performed on the culture solids retrieved from the cultivation of the fungi with wheat straw or timothy hay (Fig. 3A). Here sequential solubilization and extraction of specific cell components and quantification of the weight of residual fibres, allows quantitative analysis of the fibre content and composition of analysed plant material. Neutral detergent extraction solubilizes protein, cell content, pectins and lipids, resulting in a neutral detergent fibre (NDF) residue that corresponds to total fibre content or, broadly speaking, structural parts of plant cell walls. Subsequent acid detergent extraction removes hemicelluloses, leaving behind acid detergent fibre (ADF) consisting of cellulose and lignin [43].

**Figure 3.**
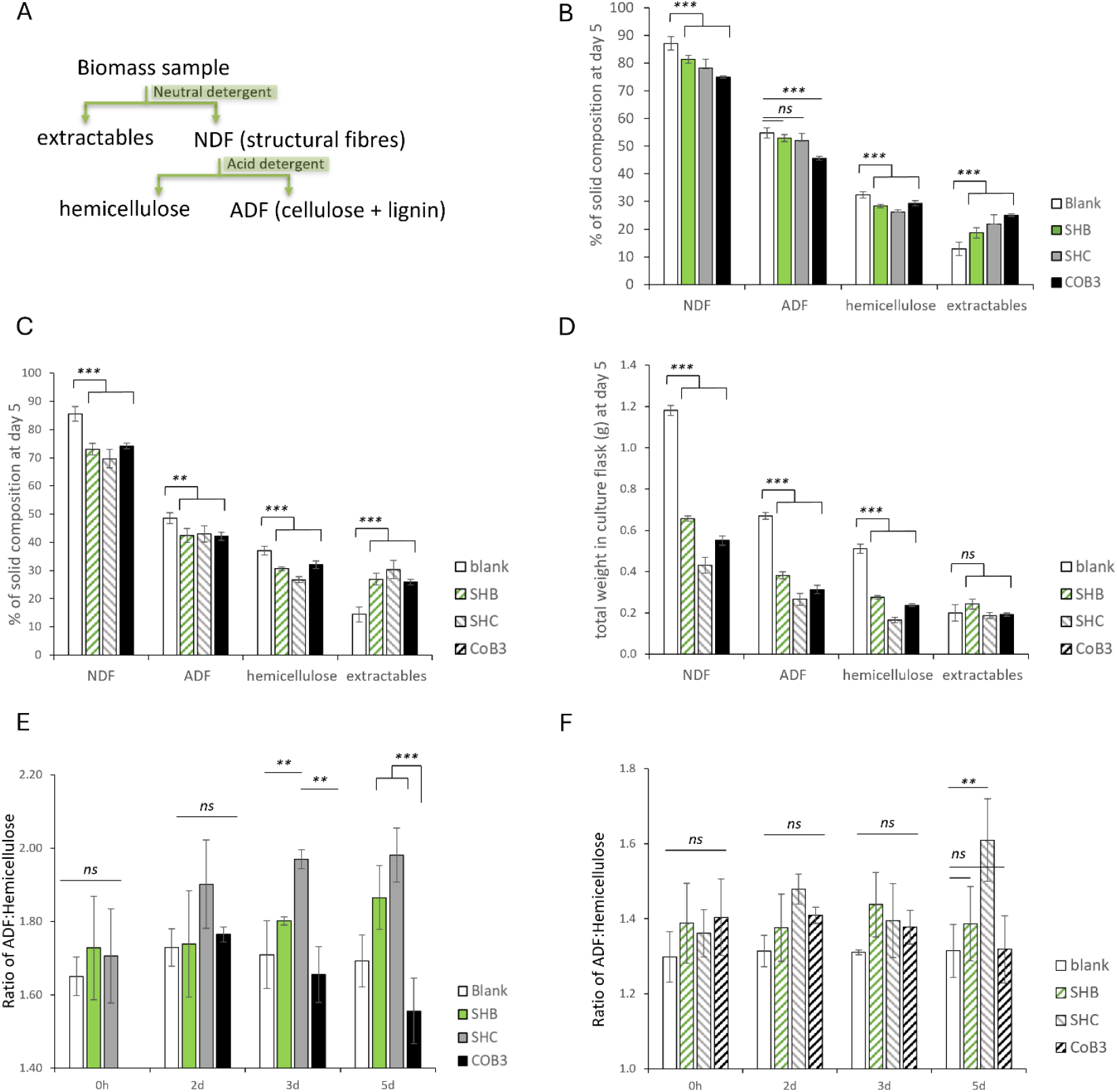
Distinct effects of AGF on lignocellulose composition. a) scheme of analysis steps. b) Composition, as % (w/w), of solids analysed from cultures with wheat straw at day 5 and c) with timothy hay at day 5. d) Total amounts of fibre and extractables per culture at day 5. e) Ratio of ADF: hemicellulose in solids from cultures containing wheat straw, and f) timothy hay. All values given as mean ± stdev (n≥3).

#### 3.3.1 Digestion efficiency of structural plant cell wall components not directly linked to fungal metabolic activity

A reduction in NDF, ADF and hemicellulose content was observed for both wheat straw (Fig. 3B) and timothy hay (Fig. 3C) culture solids. The amount of non-fibers, or extractables, found in culture solids relatively increased over time for both substrates (Fig. 3B,C), however, when analysed in absolute amounts per culture flask (Fig. 3D) it becomes clear that the total amount of extractables per culture flask stayed constant during cultivation (Fig. 3C), while NDF, ADF and hemicellulose were depleted.

For wheat straw, similar amounts of NDF eventually remained in the culture for CoB3 (51 ± 1%) and SHC (54 ± 11%), while for timothy hay (Fig 3D), a higher amount of NDF remained for CoB3 than for SHC (47 ± 2 % vs 37 ± 3 % NDF, respectively). These results may indicate differences in the fermentation efficiency between the filamentous fungal isolates SHC and CoB3, whereby *P. spp* SHC has a similar (on wheat straw) or higher (on timothy hay) ability to solubilize and digest lignocellulose than *N. frontalis* CoB3, despite the higher growth rate and metabolic activity of *N. frontalis* CoB3. For *C. communis* SHB, the in comparison high amount of NDF remaining on both wheat straw (71 ± 4 %) and timothy hay (56 ± 1 %) did coincide with the relatively low growth rate of this isolate.

#### 3.3.2 The tested fungal species each had distinct preferences for digestion of cellulose or hemicellulose

The cultivation of *N. frontalis* CoB3 with wheat straw resulted in differences in the type of fibres that were consumed from the cultures, compared to cultivation of *C. communis* SHB and *P. spp* SHC. The ratio between the total digestion of ADF (mainly comprised of cellulose + lignin) and hemicellulose in the culture (Fig. S3A) was <1 for SHB and SHC (0.79 ± 0.14 for SHB and for SHC 0.83 ± 0.11 on day 5) indicating that relatively more hemicellulose than ADF has been digested during cultivation. However, for CoB3, the ratio between the ADF and hemicellulose digestion was ≥1 (1.08 ± 0.06 at day 5), indicating that in total similar or more ADF had been digested than hemicellulose.

Similarly, the effect of the fungal activity on relative composition of the remaining plant material distinguished CoB3 from SHB and SHC (Fig 3E). The ratio between the percentage of ADF and hemicellulose remaining in the solids of wheat straw cultures of the blank control cultures is 1.70 ± 0.07, and for *N. frontalis* CoB3 no significant changes in this ratio were detected during cultivation, suggesting that the digestion of hemicellulose and ADF had taken place at similar rates, resulting in an unchanged relative composition. For *C. communis* SHB and *P. spp* SHC, the ratio increased to 1.87 ± 0.09 and 1.98 ± 0.07, respectively, and thus relatively more ADF than hemicellulose was remaining in culture solids, indicating preferential digestion of hemicellulose had taken place. This aligns with the preference of these fungi for hemicellulose consumption from the total culture calculated above. No changes were detected in the composition of material from the not inoculated control cultures over time.

The effect of the fungi on timothy hay fibers was slightly different to wheat straw. For both SHB and CoB3 the ratio of ADF to hemicellulose in remaining solids remained stable (Fig 3F). For SHC, this ratio increased over time, similar as found for samples from wheat straw cultures, indicating that only for SHC relatively more hemicellulose than ADF has been digested during cultivation on timothy hay. Consistent with this, the ratio between the total digestion of ADF and hemicellulose in the culture (Fig. S3B) was for SHC 0.90 ± 0.04 on day 5, indicating that more hemicellulose than ADF has been digested. However, for CoB3 and SHB, the ratio between the ADF and hemicellulose digestion was ~1, (1.02 ± 0.03 for CoB3 and 0.96 ± 0.09 for SHB at day 5), indicating that in total similar amounts of ADF and hemicellulose had been digested.

Similar trends can be identified from the calculation of the total digestibility of NDF (NDFD), ADF (ADFD) and that of the hemicellulose fraction (Table 1). Notably, while the digestive activity of *C. communis* SHB is more effective towards all fractions of timothy grass hay than wheat straw, the largest increase (1.7-fold) is seen in the digestion of ADF fraction. This suggests that differences in the structure of the cellulose - lignin complex between the two substrates, or the accessibility of these structures, are particularly affecting activity of this fungus.

**Table 1.**
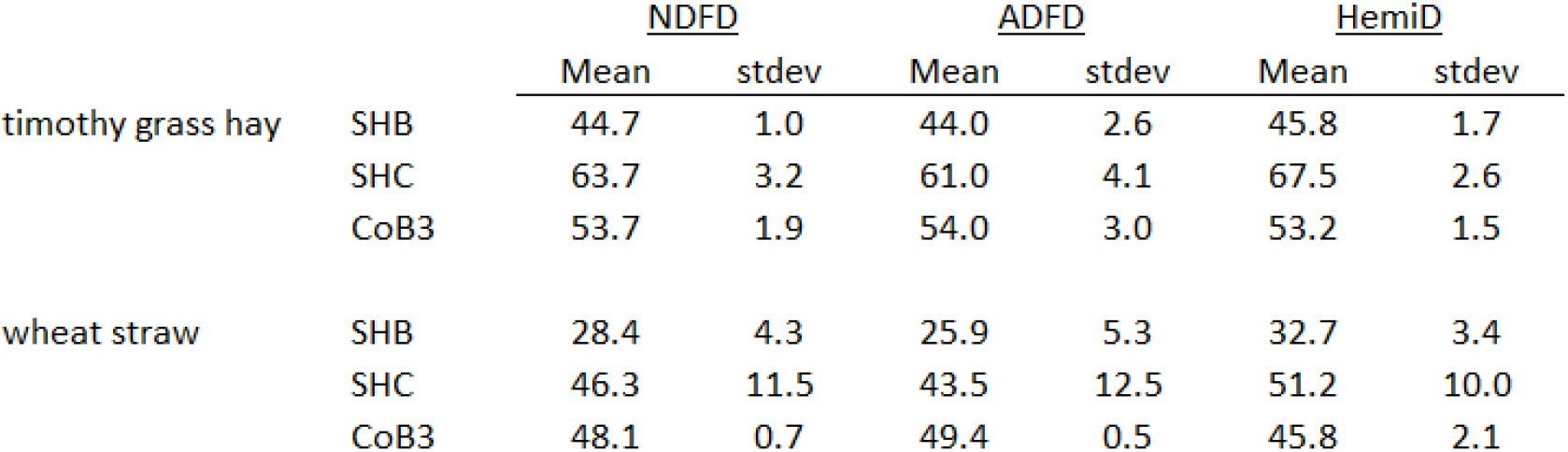
Digestibility of plant material in fungal cultures. Values, given as %, are the mean of ≥ 3 replicates. stdev, standard deviation; NDFD, Neutral detergent fibre digestibility; ADFD, Acid detergent fibre digestibility; HemiD, hemicellulose digestibility.

|  |  | NDFD |  | ADFD |  | HemiD |  |
| --- | --- | --- | --- | --- | --- | --- | --- |
|  |  | Mean | stdev | Mean | stdev | Mean | stdev |
| timothy grass hay | SHB | 44.7 | 1.0 | 44.0 | 2.6 | 45.8 | 1.7 |
|  | SHC | 63.7 | 3.2 | 61.0 | 4.1 | 67.5 | 2.6 |
|  | CoB3 | 53.7 | 1.9 | 54.0 | 3.0 | 53.2 | 1.5 |
| wheat straw | SHB | 28.4 | 4.3 | 25.9 | 5.3 | 32.7 | 3.4 |
|  | SHC | 46.3 | 11.5 | 43.5 | 12.5 | 51.2 | 10.0 |
|  | CoB3 | 48.1 | 0.7 | 49.4 | 0.5 | 45.8 | 2.1 |

### 3.4 Ability of fungi to grown on lignocellulose-derived monosaccharides

The apparent distinct fibre abilities and preferences in the three fungal isolates *N. frontalis* CoB3, *C. communis* SHB and *P. spp* SHC could be linked to their metabolic capacity for uptake and growth on degradation products from hemicelluloses or cellulose. To assess if indeed the three fungi differed in their ability to metabolise common plant-derived monosaccharides, we tested if growth of our isolates could be sustained on media containing either glucose, mannose, galactose, xylose or arabinose as carbon source. Each of the fungi accumulated fermentation gas pressure only when grown on glucose (Fig. 4A), which coincided with a reduction in culture medium pH only on this substrate (Fig. 4B), thus demonstrating that the fungi can use glucose as carbon source, but not any of the other tested sugars. The metabolic ability of these fungi towards small sugars derived from digestion of plant polysaccharides does therefore not seem to be linked to the effects of the fungi on plant material during digestion.

**Figure 4.**
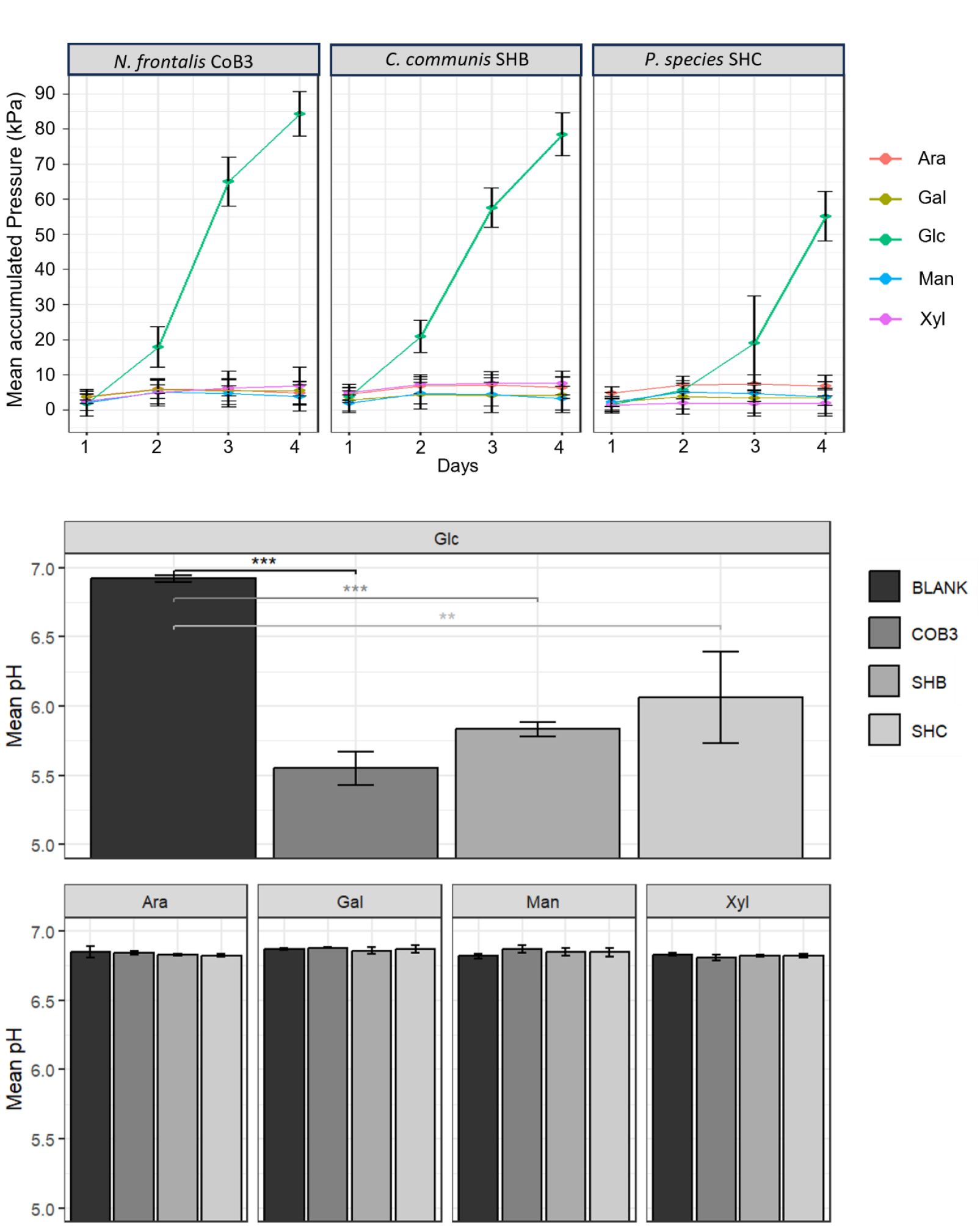
Growth of AGF on monosaccharides common in lignocellulose. a) Growth of three AGF species as detected by fermentation gas accumulation, using as carbon source arabinose (Ara), galactose (Gal), glucose (Glc), mannose (Man) or xylose (Xyl). b) pH of culture supernatant upon harvest. Values given are mean ± stdev (n=6).

## 4. Discussion

This study investigated the effects of distinct AGF species on composition and fermentation of grass and wheat straw, which are materials that as forage form part of the diet of agriculturally important ruminants, and serve as feedstock for renewables-based biotechnology processes. We hypothesized that fungi with distinct morphological properties and genetic capacity, have distinct abilities to digest plant material, and therefore different effects on the plant lignocellulose structure and composition, which overall reflects the fungal distinct niches. In summary, our results confirm that three anaerobic fungi commonly found in agriculturally important ruminants, *N. frontalis, C. communis* and *P. spp*, have different effects on plant lignocellulose properties during its digestion, which suggests that these fungi have different niches in the digestive system.

Knowledge about the specific niches that AGF occupy, or the fungal traits that enable this occupation, is very limited. Distinct distribution of the currently 88 recognised genera of AGF in herbivorous animals has been reported, with strong association of some AGF genera with certain animals, for example a *Khoyollomyces* with horses and zebras, and the so far uncultured genus NY20 to rhinoceroses [11]. This strongly suggests links between traits of these fungi and the niche available in the environment provided by the animal host, influenced by factors such as feed composition, animal physiological parameters or wider microbiome context. However, the phylogeny of the 28S and ITS regions of the distinct fungi that we isolated for this study demonstrates that the AGF isolates are robustly placed in genera *Neocallimastix, Caecomyces* and *Piromyces*. In a recent expansive study of the herbivore mycobiome [11], the AGF genera studied here were present in the top 8 of most abundant AGF genera, with *Piromyces* at 10%, *Caecomyces* at 6% and *Neocallimastix* at 5%, and, as reviewed previously [37], were found in a wide range of hosts. Therefore, existence of any specific niches, or fungal traits enabling occupation of such niches, have so far been harder to derive for these genera compared to more specialised AGF genera.

The results from this study provide an initial insight in the potential niches of species of genera *Neocallimastix, Caecomyces* and *Piromyces* in relation to lignocellulose digestion. We found that the distinct fungi have different effects on plant material during its digestion: *N. frontalis* CoB3 digested the hemicellulose and cellulose containing fractions of the wheat straw and timothy hay with equal efficiency, while *P. spp* SHC preferentially digests hemicellulose in both substrates, and was more effective growing on and digesting the substrate high in hemicellulose content. Meanwhile, the result of activity of *C. communis* SHB on the composition of the plant material differed per substrate, with preferential hemicellulose consumption on wheat straw but not on timothy hay. *C. communis* SHB was able to degrade timothy grass hay, and particularly its cellulose and lignin fraction, much more effectively than wheat straw, while *N. frontalis* CoB3 seems adapted to degrade both substrates almost equally well.

Previous work of Theodorou and colleagues assessed the effects of *Neocallimastix* sp strain R1 on the composition of Italian ryegrass hay [44], whereby the hay used as carbon source in these cultures was consumed to the same extend (47-55% reduction in dry weight) as in this study. The monosaccharides arabinose, galactose, glucose and xylose were removed from the rye grass concurrently and at similar rates during the main phase of biomass removal [44]. Also in perennial ryegrass, the final extent of removal of glucose and xylose by *Neocallimastix* sp. strain CS3b has been reported to be similar [45], and as these monosaccharides are the main building blocks of cellulose and the most common hemicellulose xylan, respectively, this suggest that also here *Neocallimastix* sp. degradation cellulose and hemicellulose at a similar rate. These findings are similar to the concurrent digestion of hemicellulose and cellulose that we report here for *N. frontalis* on both analysed plant substrates, suggesting this aspect of the *Neocallimastix* digestion strategy may be conserved across different strains and across complex substrates.

Various studies have investigated the effect of AGF on digestibility of wheat straw and other ruminant feed components but often don’t report results from axenic cultures nor the content or digestibility of hemicelluloses, making it more difficult to compare the outcomes of these studies to the work presented here. However, treatment of plant materials with AGF, particularly *Neocallimastix* or *Piromyces* species have been reported to reduce NDF and ADF content and therefore increase digestibility, and this is consistent with our findings. *Piromyces* Yak-G18 cocultures with *Methanobrevibacter ruminantium* reached NDFD of 41% and ADFD of 31% after 5 days incubation on wheat straw [46]. The reported higher NDFD suggests a considerably higher digestibility of the hemicellulose fraction compared to the cellulose and lignin cell wall fraction, similar as found in this study for activity of *Piromyces sp*. A *N. frontalis* YakQH5 coculture with the methanogenic archaea *Methanobrevibacter gottschalkii* on wheat straw for 5 days resulted in 49.5% NDFD, similar to the 48% NDFD identified here for the *N frontalis* CoB3 axenic culture [47]. A *N. frontalis* Yaktz1 axenic culture achieved a NDFD of ~ 40% and ADFD of ~ 35%, both considerably lower than found here for *N. frontalis* CoB3. [48]. Culture conditions or fungal strain differences may play a role in this discrepancy and highlights the importance of comparing results of fungal activity on plant material composition under comparable conditions.

*Caecomyces* may occupy a distinct niche from *Neocallimastix* and *Piromyces*. Our results indicate that *C. communis* isolate SHB was much more effective in digesting timothy grass hay than wheat straw, especially its cellulose and lignin containing fraction (1.7-fold increase in digestibility), whereas *N. frontalis* CoB3 digested both substrates with similar efficiency. Botten *et al* reported that a *Caecomyces* isolate colonised specific cell types, lignified secondary xylem fibres, in xylem of alfalfa hay, while *Neocallimastix* and *Piromyces* isolates colonised all cell types [49]. Both these findings suggests that Caecomyces species may be more specialised than *Neocallimastix* and *Piromyces* species with regard to the degradation and use of lignocellulose components. It has also been suggested that *Caecomyces* species may occupy a niche with higher reliance on soluble sugars compared to other AGF. *C. churrovis* is reported to transcribe a higher proportion of soluble CAZymes compared to those in multi-enzyme complexes (fungal cellulosomes), when comparing to *N. californiae, Anaeromyces robustus* and *P. finnis* [50]. *Caecomyces* are more often found in faecal samples compared to rumen content samples, which has been suggested to possibly link to their increased reliance on soluble carbohydrates, substrates mostly available in the colon [13]. We found the *C*.*communis* isolate SHB is in comparison to *N. frontalis* and *P. spp*. less effective in all tested measures of plant degradative activity, it has a relatively low growth rate and the most limited digestion of complex biomass in terms of dry weight, NDFD and ADFD. This aligns with low growth rates reported for other *Caecomyces* isolates growing on complex plant material. For example, the growth rates of 0.028 to 0.039 h^−1^ for *C. churrovis* on reed canary grass and corn stover [50] are similar as found here for *C. communis* SHB with 0.031 to 0.038 h^−1^ on timothy hay and wheat straw. However, [51] found that during ex vivo fermentations of complex plant biomass by the bovine rumen microbiome, supplementation of the rumen fluid with a *Caecomyces* isolate was more effective than supplementation with other AGF with regard to fermentation effectivity on complex plant material. Together this suggests that the *C. communis* SHB low growth rate and fermentation effectivity found in this study may not linked to the used growth conditions. Instead, as further suggested by the fact that this genus is a stable member common in ruminants mycobiomes [13], [20] it is likely that other properties or traits provide benefits to enable species from this genus to be competitive with other AGF and achieve stable niche occupation. These could be traits regarding interaction with other rumen microbes, gut physiology or resilience under stress conditions.

The consumption of plant-derived monosaccharides was for each of the AGF isolates limited to glucose. Many AGF have been reported to have a limited use of plant-derived monosaccharides, with notable exceptions such as *Feramyces austinii* [52] and *Aestipascuomyces dupliciliberans* [39]. However, for *Piromyces* and *Neocallimastix* isolates, genes encoding the full xylose isomerase pathway have been identified [53], [54], and activity of the two enzymes of this pathway, xylose isomerase and D-xylulokinase was demonstrated in crude extracts of *Piromyces sp*. strain E2 (Harhangi et al., 2003). Despite this, *N. lanati* and *N*.*californiae* were reported not to grow on xylose [54], [55], similar to what we found here. Differences in uptake conditions, transporters and other so far unidentified details of growth conditions may be responsible for these observed discrepancies. For *C. communis* var. *churrovis*, gene annotations for the kinase enzyme of the xylose catabolic pathway were reported to have low confidence and the lack of growth of this species on xylose, which is similar to what we observe here for *C. communis* SHB, was suggested to maybe due to an incomplete metabolic pathway (Henske et al., 2017).

AGF species have been demonstrated to increase the expression of expansive sets of genes encoding plant cell wall degradative enzymes during exposure to complex plant material, for example [50], [56], [57]. While regulation underlying such transcriptomic responses is still unknown, these studies confirm that AGF have the machinery to digest a range of plant polysaccharides, even those that would result in degradation products, such as xylose and mannose, that they do not seem to be using for metabolism or growth. Our analysis of plant material indicates that the three investigated AGF digest both hemicelluloses and cellulose cell wall fractions, while having different plant biomass degradation strategies. A side-by-side comparison of the transcriptome of these different species may elucidate the fungal activity responsible for the observed effects of their distinct degradation strategies.

## Ethics

Rumen fluid was collected adhering to the guidelines and regulations of the UK Home Office Animals (Scientific Procedures) Act of 1986, with animal maintenance and sampling covered by Home Office project licence PP7153972. All experimental protocols were approved by the project licence holder and the University of Glasgow Animal Welfare and Ethical Review Board and adhere to ARRIVE guidelines.

## Data accessibility

The ITS and 28S gene fragments sequenced for this study are accessible in NCBI via accession number PX270339-PX270347 and PX270323-PX270334. Supplementary material is available online.

## Declaration of AI use

We have not used AI-assisted technologies in creating this article.

## Authors’ contributions

JvM conceived the studies and designed experiments, SS JLM SL and JvM performed experimental work, SS, JLM and JvM performed data analysis. JvM secured funding and supervised the project. SS, JLM and JvM generated figures and wrote the manuscript, all authors reviewed the final manuscript and agreed with its content.

## Acknowledgements

We are grateful to farm staff for enabling rumen fluid collections. We thank Eva Ramos-Morales for practical advice when starting anaerobic cultivations, Neil Graham for donating wheat straw and John Parker for help with milling the straw.

---

**Figure S1.**
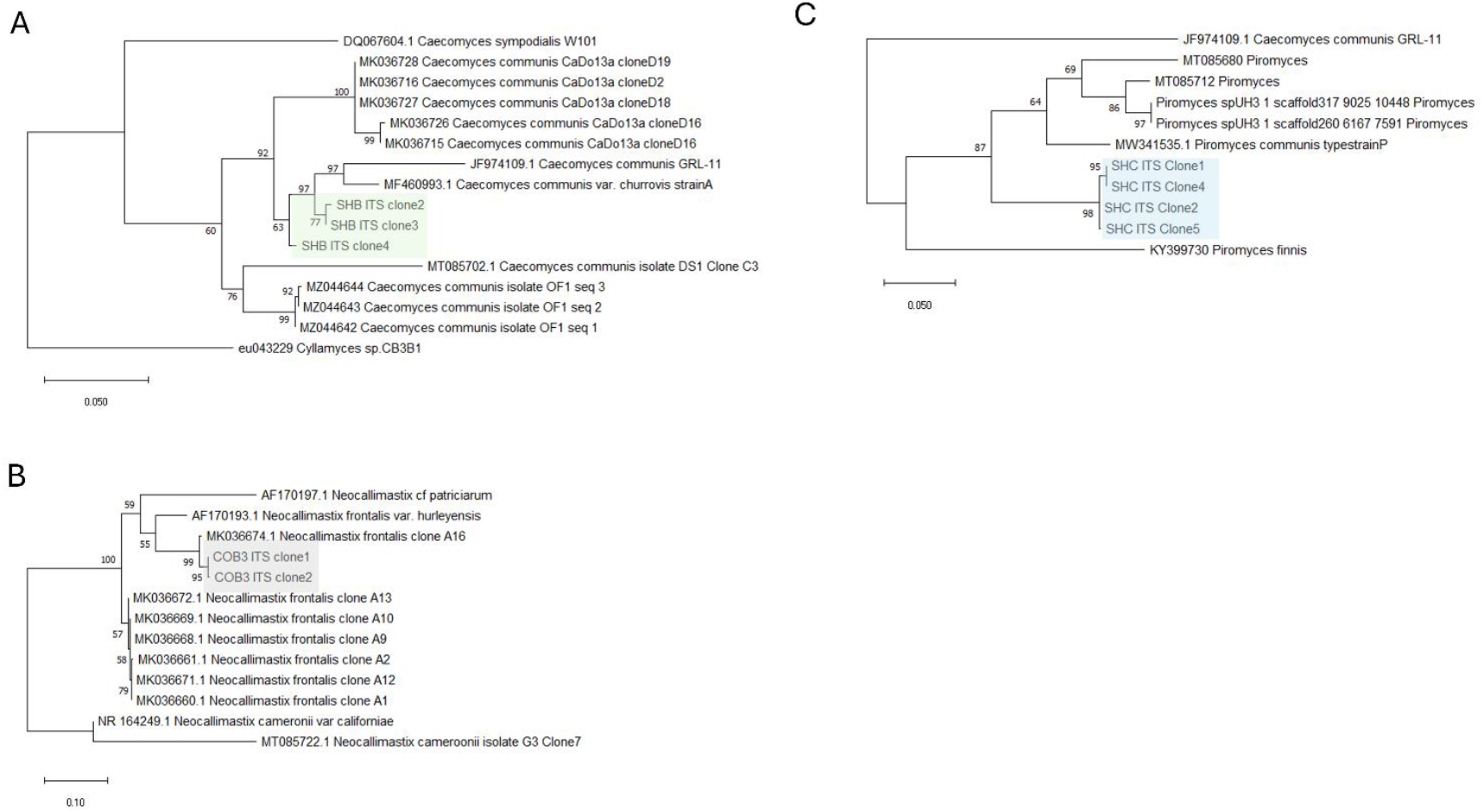
Phylogeny of three AGF isolates based on maximum likelihood tree of ribosomal internal spacer region (ITS) DNA sequences. With a) *N. frontalis* CoB3, b) *C. communis* SHB, c) *P. spp* SHC.

**Figure S2.**
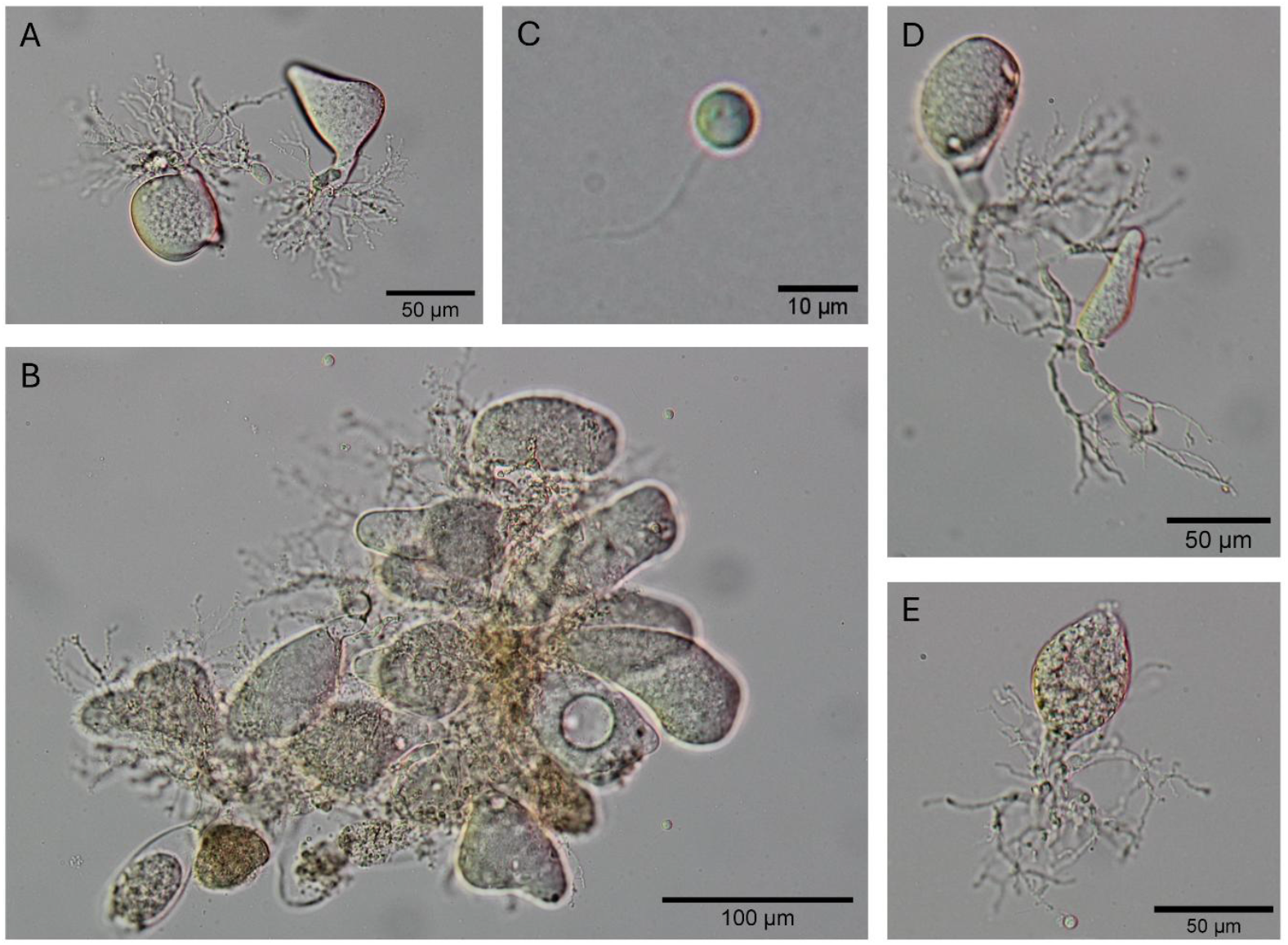
Additional images of *P. spp* SHC. With a, b) thalli showing irregular morphology including triangular and heart-shaped c) uniflagellar zoospore d,e) short, thin and heavily branched rhizoidal morphology. All images taken from 2 day old cultures grown on media C with glucose as carbon source.

**Figure S3.**
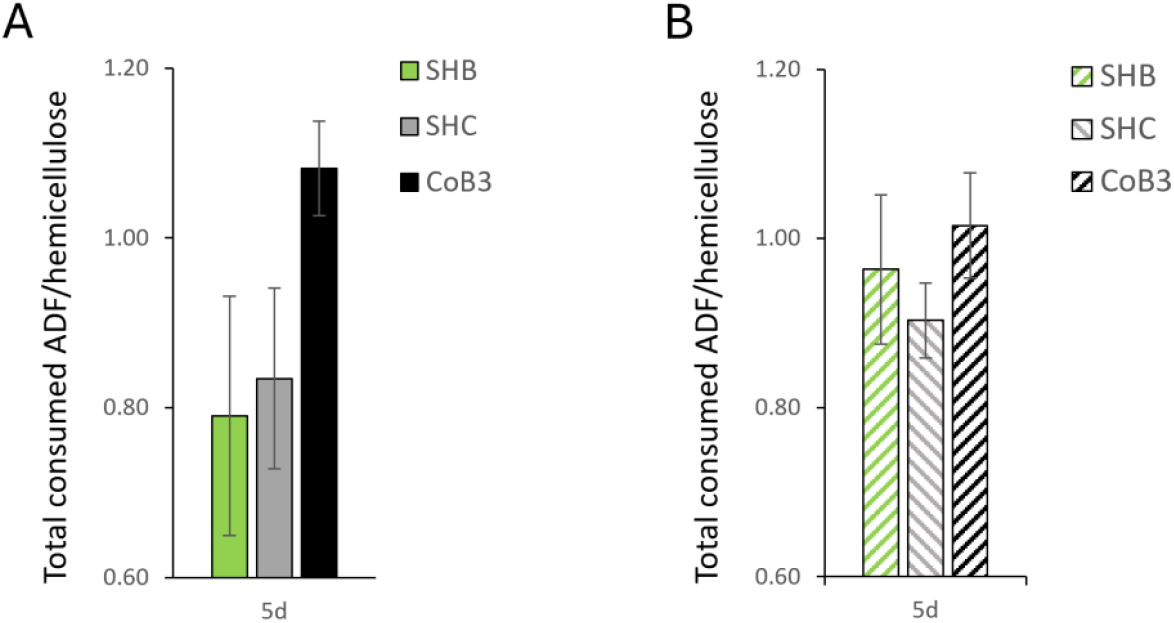
Ratio between the total digestion of ADF and hemicellulose fractions in the culture flasks. a) in cultures containing wheat straw and b) in cultures containing timothy grass hay. Values are the mean (n ≥ 3) ± stdev

